# Cumulative timing-dependent changes in corticospinal excitability during suprathreshold paired-pulse transcranial magnetic stimulation

**DOI:** 10.1101/2025.11.19.689398

**Authors:** Suraj Suresh, Mana Biabani, Wei-Yeh Liao, George M Opie, Alex Fornito, Mitchell R Goldsworthy, Nigel C Rogasch

## Abstract

Transcranial magnetic stimulation (TMS) is widely used to assess inhibitory and facilitatory circuits within the primary motor cortex. However, accumulating evidence suggests that even brief TMS paradigms may induce unintended changes in corticospinal excitability. Here, we examined whether suprathreshold paired-pulse TMS delivered at inter-stimulus intervals (ISIs) associated with intracortical facilitation or long-interval cortical inhibition (LICI) elicits cumulative changes in motor-evoked potential (MEP) amplitude. In experiment 1, we reanalysed data from 17 participants who received 20 suprathreshold paired-pulses at eight ISIs (10–200 ms) and 40 unconditioned single pulses. Stimulation was pseudo-randomised and distributed evenly across four blocks. A linear mixed-effects model assessed trial-wise changes in MEP amplitude across ISIs. We replicated the design in an independent sample (n=10, experiment 2). A significant trial-by-ISI interaction was observed in both cohorts. Specifically, MEP amplitudes increased across trials for ISIs of 20 and 30 ms (p<0.05), but remained stable at LICI-related ISIs (100–150 ms). A similar increase was also seen with single-pulse TMS. These findings demonstrate that suprathreshold paired-pulse TMS at short ISIs can cumulatively enhance corticospinal excitability during stimulation. Furthermore, the results suggest potential for using these protocols not just for probing cortical circuits, but also as interventions to modulate motor system excitability.

## 1. INTRODUCTION

Transcranial magnetic stimulation (TMS) is a powerful tool to non-invasively probe excitatory and inhibitory circuits within the human cortex [1]. Above a certain threshold, applying a single TMS pulse to the primary motor cortex results in a compound muscle action potential in peripheral muscles controlled by the targeted cortical region. These responses can be measured using electromyography (EMG) and are known as a motor-evoked potential (MEP) [2]. The amplitude of the MEP provides a simple measure of the excitability of the corticospinal system and is often used to track changes in excitability during different brain states, in brain disorders, and following neuromodulatory TMS paradigms like repetitive TMS (rTMS) [2-4].

In addition to single-pulse TMS, paired-pulse TMS is a methodology used to assess activity in inhibitory and excitatory intracortical circuits [5]. Paired-pulse stimulation involves delivering a conditioning stimulus and a test stimulus, separated by an interstimulus interval (ISI) [6]. By adjusting the ISI and stimulus intensities, paired-pulse stimulation is able to target different intracortical circuits. For example, using a subthreshold conditioning stimulus, MEP amplitudes following the test stimulus are reduced at ISIs between 1-6 ms, an effect referred to as short-interval cortical inhibition (SICI). In contrast, test MEPs are increased at ISIs between 10-15 ms, which is known as intracortical facilitation (ICF) [6-10]. Pharmacological studies have demonstrated that SICI is mediated by type A gamma-aminobutyric acid (GABA_A_) receptors, whereas ICF is potentially mediated by excitatory n-methyl d-aspartate (NMDA) receptors [11-13]. When a suprathreshold conditioning pulse is used, MEP amplitudes are increased at ISIs between 10-40 ms, but reduced at longer ISIs between 50-150 ms. While the mechanisms of facilitation at long-intervals remain unclear – possibly involving both spinal and cortical processes [14, 15] – inhibition at long-intervals (i.e., long-interval cortical inhibition; LICI) is mediated by type B GABA (GABA_B_) receptors [12, 13, 16]. Both single- and paired-pulse TMS paradigms have provided valuable insights into the neurophysiology of healthy and pathological brain function[5, 17].

An underlying assumption of using single- and paired-pulse TMS paradigms to investigate cortical excitability and inhibition is that the delivered pulses do not modulate excitability (i.e., induce neural plasticity) [2, 5]. MEP amplitudes naturally vary from trial to trial. As a result, it is common practice to record between 5-20 trials for each single/paired pulse condition investigated and then average across trials to quantify corticospinal excitability [18]. This can result in hundreds of stimuli within an experimental session. To avoid modulating cortical excitability by inducing neural plasticity, single- and paired-pulse paradigms are typically delivered with an interval of at least 3 seconds between trials and a jittered inter-trial interval [1, 17]. However, several studies have demonstrated that MEPs following single-pulse TMS at intervals of 3-6 seconds show cumulative increases in MEP amplitude across trials, consistent with the induction of neural plasticity [19, 20]. Similar cumulative changes in excitability have been reported using paired-pulse paradigms with ISIs targeting excitatory circuits and subthreshold stimuli [21, 22]. However, it remains unclear whether similar cumulative changes in corticospinal excitability occur with suprathreshold paired-pulse paradigms with longer ISIs, like those targeting suprathreshold facilitation and LICI.

The aim of this study was to assess whether conditioned MEP amplitudes are modulated across trials using suprathreshold paired-pulse paradigms with longer ISIs (≥10 ms). First, we performed an exploratory analysis on an existing dataset which collected MEPs following suprathreshold paired-pulses from eight ISIs between 10-200 ms [22]. We then collected a second dataset using the same protocol as the initial study to test the replicability of the findings.

## 2. METHODS

### 2.1. Participants

This study analysed data from two single-session experiments that included paired-pulse TMS applied at suprathreshold intensities. In experiment 1, we performed an exploratory analysis on existing data from 17 participants (25 ± 6 years; 11 females) previously collected and published by our group at Monash University (human ethics reference number: 6054) [23]. While the original publication only included 16 participants in their final analysis, reasons for exclusion were unrelated to the data used in this study. The current study was therefore able to include data from all 17 participants. In experiment 2, we replicated experiment 1 in an independent cohort of 10 participants (31 ± 4 years; 6 females) to validate the findings from the exploratory analysis. Data were collected at the University of Adelaide (human ethics reference number: H-2021-029). Before performing experimental procedures, informed consent, demographic information, a handedness questionnaire and TMS safety compliance were obtained from all participants [23-25].

### 2.2. Experimental design

Experiment 2 was modelled after Experiment 1 (see Biabani et al., 2021) with minor modifications. The experiment commenced with a preparation phase to determine the hotspot position for left primary motor cortex and two baseline measures to determine stimulation intensity (resting motor threshold [RMT] and 1 mV threshold [S1mV]). Single- and paired-TMS pulses were then delivered at 0.2 Hz (±1 second jitter) to the left primary motor cortex. Single-pulses were delivered at S1mV, whilst paired-pulses were delivered with the conditioning pulse at 120% RMT and the test pulse at S1mV. Eight different ISIs were used for paired-pulse stimulation (10, 20, 30, 40, 50, 100, 150, and 200 ms). The experiments consisted of four 50 trial blocks, each consisting of 10 single-pulses and 40 paired-pulses. The eight paired-pulse conditions were evenly split across each block, resulting in five repetitions per ISI within each block. The order of single- and paired-pulse trials were pseudo-randomised within each block.

### 2.3. Electromyography (EMG)

Both experiments collected EMG from two Ag-AgCl surface electrodes over the right first dorsal interosseous (FDI) muscle positioned in a belly-tendon montage approximately 2 cm apart. The EMG signals were band-pass filtered (10-1000 Hz), amplified (×1000), and digitized at a sampling rate of 5 kHz. EMG data was processed offline, using Lab Chart 8 (ADInstruments, Sydney, Australia) in Experiment 1 and Signal software (V6) for Experiment 2. MEP peak-to-peak amplitude served as the primary outcome measure for statistical analyses.

### 2.4. Transcranial Magnetic Stimulation (TMS)

For both experiments, TMS was delivered via a figure-of-eight coil (D70, 70 mm diameter) connected to two Magstim 200^2^ stimulators (Magstim Company, Whitland, UK) via a BiStim2 module. In experiment 1, coil position was maintained using a participant-specific neuro-navigation system (Brainsight™ 2, Rogue Research Inc., Canada). In experiment 2, coil position was maintained using a mark drawn on the participant’s head using a felt tip pen. Both experiments used Signal software (V6) to deliver TMS.

The optimal left primary motor cortex hotspot was identified as the coil position that consistently elicited the largest peak-to-peak MEP amplitude from the FDI muscle. RMT was determined by gradually adjusting the percentage of maximum stimulator output (MSO%) to find the minimum intensity that produced MEPs ≥50 µV in at least five out of 10 trials. The S1mV was determined by identifying the lowest MSO% that produced MEPs with a mean amplitude of ∼1 mV across 20 single-pulse trials.

### 2.5. Data Processing

EMG data were epoched from -200 ms to 500 ms relative to the test pulse and processed offline using Lab Chart 8 (ADInstruments, Sydney, Australia) for experiment 1 and Signal software for experiment The MEP amplitude following the test stimulus were quantified for each trial by measuring the maximum peak-to-peak amplitude between 20-40 ms post the test stimulus in the respective software packages. Additionally, any trial recording which captured a peak-to-peak amplitude MEP amplitudes >25 μV prior to the TMS pulse artifacts were excluded from the analysis and were subsequently categorized according to condition (single-pulse and eight paired-pulse conditions).

### 2.6. Statistics

#### 2.6.1. Linear mixed effects model (LMM)

R statistical software (R 4.4.2 + RStudio 202.12.0) was used to fit linear mixed effects models (*lme4* package [26]) and generate *post-hoc* comparisons (*emmeans* package; [27]). Two models were specified: model 1 used data from experiment 1 only, whereas model 2 included data from both experiments 1 and 2 to facilitate comparisons between experiments. In both models, the dependent variable was single-trial conditioned MEP amplitude following paired-pulse TMS, with random intercepts for participants. Model 1 assessed the main effects of ISI (10, 20, 30, 40, 50, 100, 150, and 200 ms) and trial (trials 1–20), as well as their interaction. Model 2 extended this framework by additionally including cohort (experiment 1 vs. experiment 2) as a fixed effect. Estimated marginal means (with 95% confidence intervals, 95% CI) were generated to compare MEP amplitude within and between ISIs, trials, and cohorts. Furthermore, estimated linear trends (with 95% CI; *emtrend* function) were generated to quantify changes in MEP amplitude across trials (1–20), and to assess whether these trends differed by ISIs or cohorts. All post-hoc analyses were adjusted using Bonferroni correction to account for type-I error. Comparisons are presented as estimated mean or slope differences with 95% CIs, providing an unstandardized estimate of effect size. Equivalent models were also constructed to assess the impact of trial (model 1 and 2) and cohort (model 2 only) on MEP amplitude following single-pulse stimulation.

## 3. Results

### 3.1. Experiment 1: exploratory analysis

Figure 1 shows MEPs from a representative participant following paired-pulse TMS with different ISIs overlayed from the mean of the first and last 3 trials. Increases in the conditioned MEP amplitude over trials are evident for the 20 and 30 ms ISI conditions. In the group data, there were significant main effects of ISI [X^2^_(7, 2522)_ =1945.59, *p* <0.001] and trial [X^2^_(1, 2522)_ = 56.23, *p* < 0.001]) on MEP amplitude, and a significant ISI x trial interaction (X^2^_(7, 2522)_ =26.60, p <0.001). Post hoc analysis revealed significant differences in mean MEP amplitude across all ISIs (*p*-value = 0.021 to <0.001), excluding contrasts between 50 vs 150 ms (*p* = 0.78), 50 vs 200 ms (*p* = 1.00), 100 vs 150 ms (*p* = 1.00) and 150 vs 200 ms (*p* = 0.05). MEP facilitation was evident at ISIs between 10-30 ms and inhibition between 50-200 ms.

**Figure 1:**
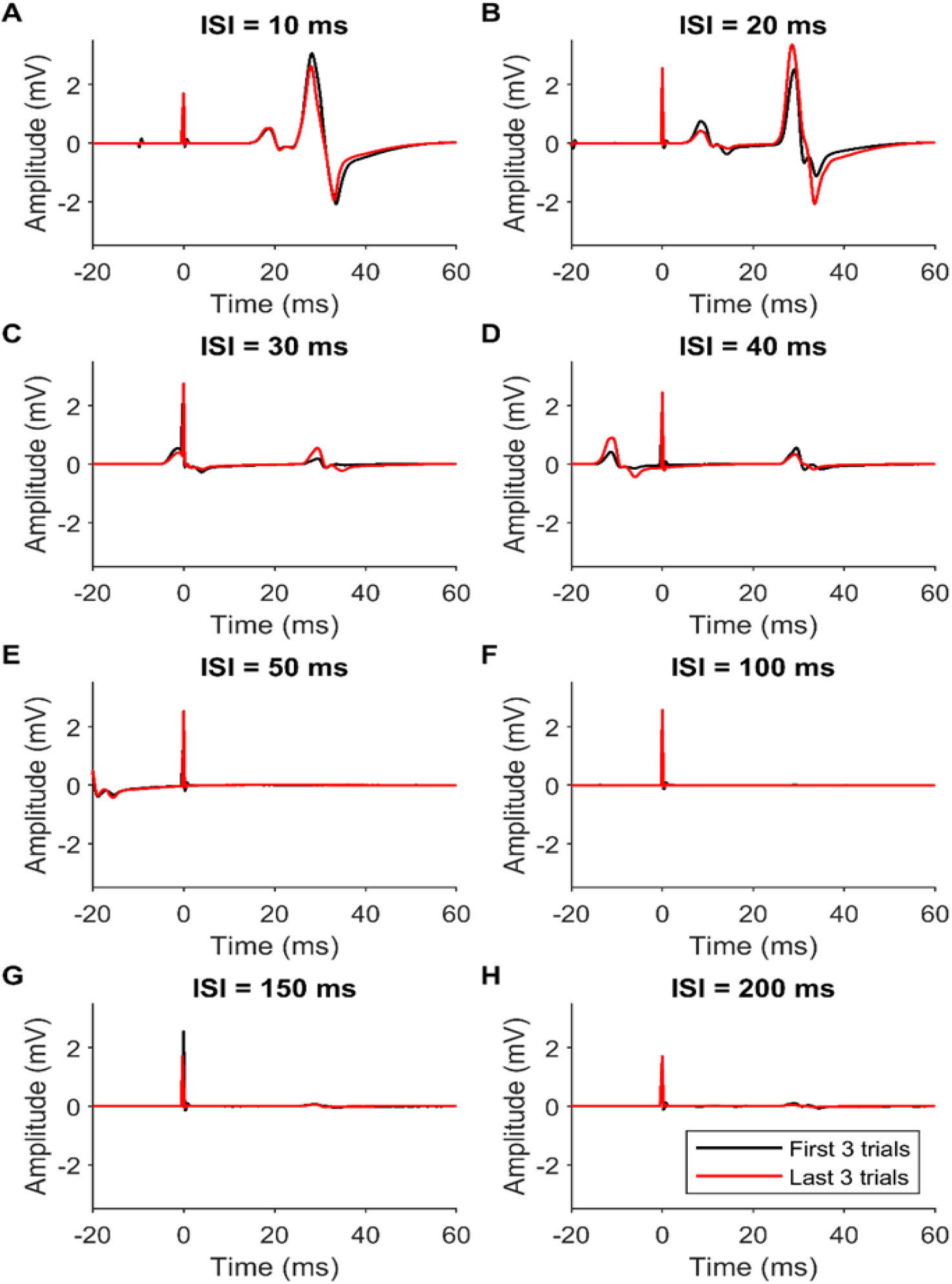
Conditioned motor evoked potentials (MEPs) following suprathreshold paired-pulse TMS at different interstimulus intervals (ISIs) from a representative participant. Traces represent the mean electromyographic signals of the first 3 trials (indicated in black) and last 3 trials (indicated in red) for each of 8 different ISIs. The test pulse was delivered at 0 ms and the latency of the conditioned MEP begins at approximately 23 ms. Increases in amplitude of the conditioned MEP in the last compared to the first 3 trials are clear in the 20 and 30 ms ISI conditions.

When comparing the changes in MEP amplitudes across trials (i.e., the slope), post hoc tests indicated significant increases in MEP amplitude over trials at ISIs of 10 ms (estimated trend = 0.020 [0.0012, 0.0379], *p* = 0.037), 20 ms (estimated trend = 0.057 [0.038, 0.075], *p* < 0.001), 30 ms (estimated trend = 0.046 [0.028, 0.065], *p* < 0.001), 40 ms (estimated trend = 0.026 [0.008, 0.045], *p* = 0.005) and 200 ms (estimated trend = 0.026 [0.007, 0.044], *p* = 0.006) (figure 2). The post hoc test contrasting MEP slopes between ISIs demonstrated significant difference in slopes between ISIs 20 vs 50 ms (*p* = 0.031), 20 vs 100 ms (*p* = 0.01) and 20 vs 150 ms (*p* = 0.002), as well as 30 vs 150 ms (*p* = 0.035), indicating that the change in MEP amplitude across trials was greater for some ISIs compared to others. Together, these results suggest that there are cumulative changes in corticospinal excitability across trials in conditions using suprathreshold paired-pulse TMS.

**Figure 2:**
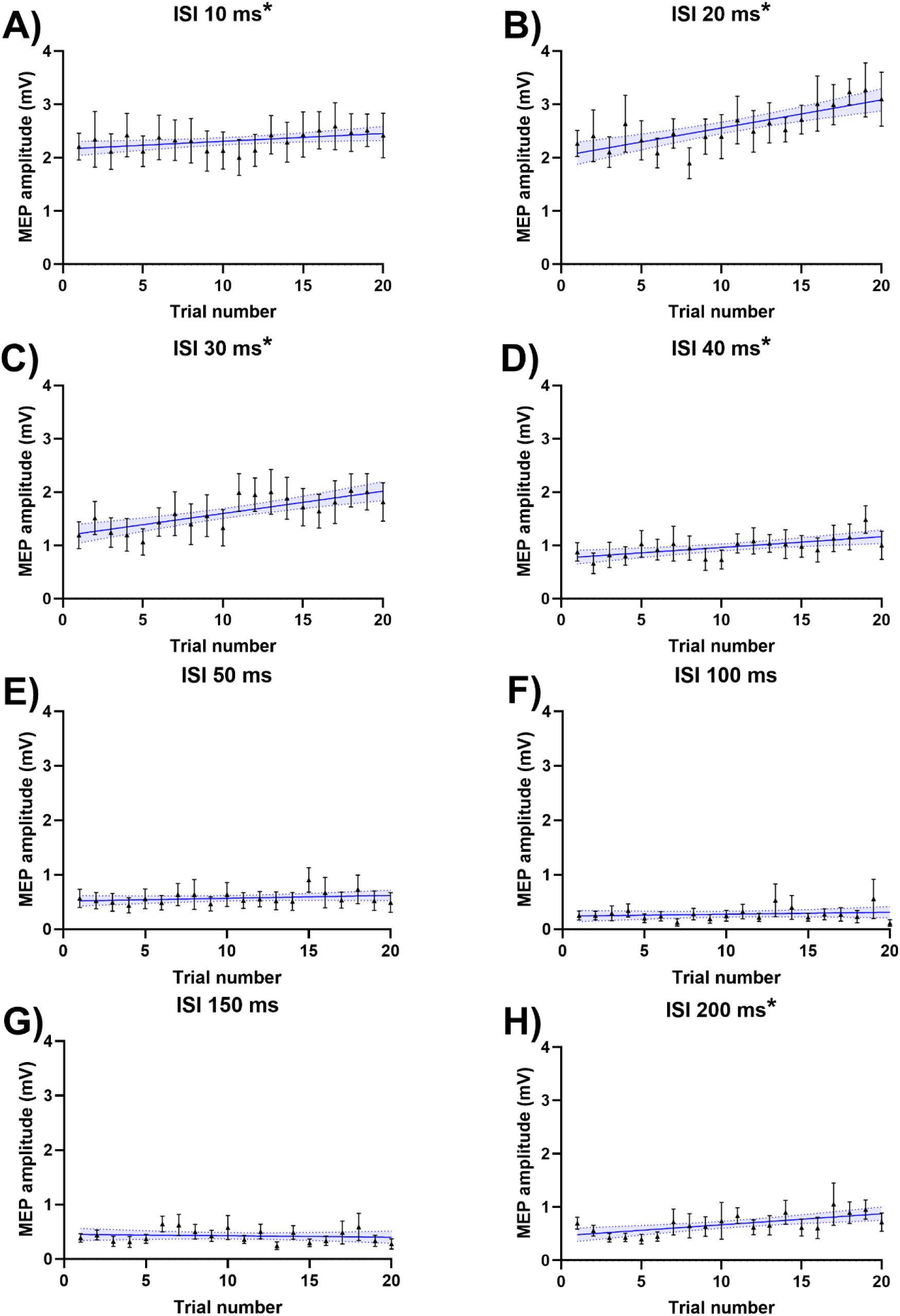
Changes in conditioned motor-evoked potential (MEP) amplitude across trials for different interstimulus intervals (ISIs) from experiment 1 (n=17). Circles indicate the mean (95% confidence intervals) conditioned MEP amplitude across participants for each trial. The blue solid line indicates the mean linear regression across trials whereas the shaded bar indicates the standard error of mean. Responses are shown for eight different ISIs including 10 ms**\*** (A), 20 ms**\*** (B), 30 ms**\*** (C), 40 ms**\*** (D), 50 ms (E), 100 ms (F), 150 ms (G) and 200 ms**\*** (H). ISIs where the linear regression slope was significantly different from 0 are indicated by an asterisk (P<0.05).

Figure 3 shows the MEP amplitude across all trials during suprathreshold single pulse delivery in experiment 1. In the group data, there was a significant main effect of trial [X^2^ _(1, 626)_ = 13.71, p = 0.0002] on MEP amplitude. To quantify the nature of this effect, a *post hoc* analysis compared the slope of the MEP curve to ‘0’ (i.e., no change); results indicated a significant positive relationship (estimated trend = 0.009 [0.004, 0.013], *P* <0.001). These results indicate that the repetitive administration of 40 suprathreshold single-pulse TMS, delivered intermittently during PPS blocks, evokes a cumulative facilitatory effect on corticospinal excitability.

**Figure 3:**
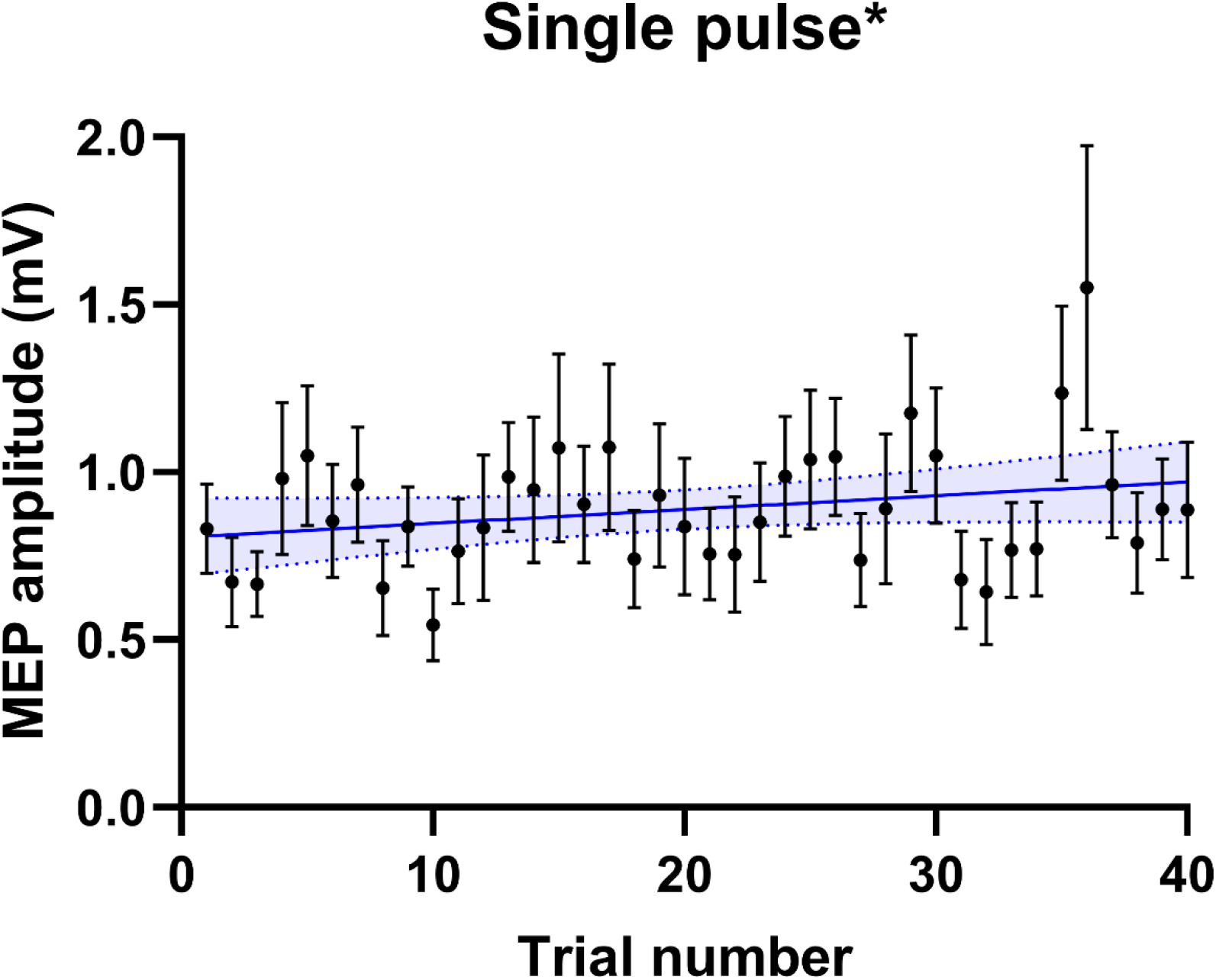
Changes in motor-evoked potential (MEP) amplitude across single pulse trials from experiment 1 (n=17). Circles indicate the mean (±SEM) MEP amplitude across participants for each trial. The blue line indicates the mean linear regression across trials whereas the shaded bar indicates 95% confidence intervals. The linear regression slope of the single-pulse trials was demonstrated a significant positive deviation from 0 (p < 0.001).

### 3.2. Comparing experiment 1 and 2

To assess the reproducibility of the main findings, a separate model was performed to contrast conditioned MEP amplitude between the two experimental cohorts. We found significant main effects of cohort (X^2^ _(1, 4105)_ = 52.452, < 0.001), ISI (X^2^ _(7, 4102)_ = 2455.6, *p* < 0.0001) and trial (X^2^ _(1, 4103)_ = 28.7, *p* < 0.001) on conditioned MEP amplitude, as well as a significant interaction between ISI and trial (X^2^ _(7, 4102)_ = 39.4, *p* < 0.001), and cohort and ISI (X^2^ _(7, 4102)_ = 26.3, *p* <0.001). Post-hoc analyses on the interaction between cohort and ISI indicated that conditioned MEP amplitudes were significantly larger for ISIs between 10–50 ms in experiment 2 compared to experiment 1 (all *p* < 0.004). However, there was no evidence for a significant interaction between cohort and trial (X^2^ _(1, 4102)_ = 2.7, *p* = 0.10), or cohort, ISI and trial (X^2^ _(7, 4102)_ = 5.7, *p*= 0.57). Furthermore, when comparing changes in MEP amplitudes across trials, we could not find evidence for a difference in slope between cohorts for ISIs between 10-150 ms, except for ISI 200 ms (EMD= 0.038, SE=0.0177, *p* = 0.03) (figure 4). Together, these findings suggest that cumulative changes in corticospinal excitability during suprathreshold paired-pulse paradigms are a robust finding that replicated across two independent cohorts.

**Figure 4:**
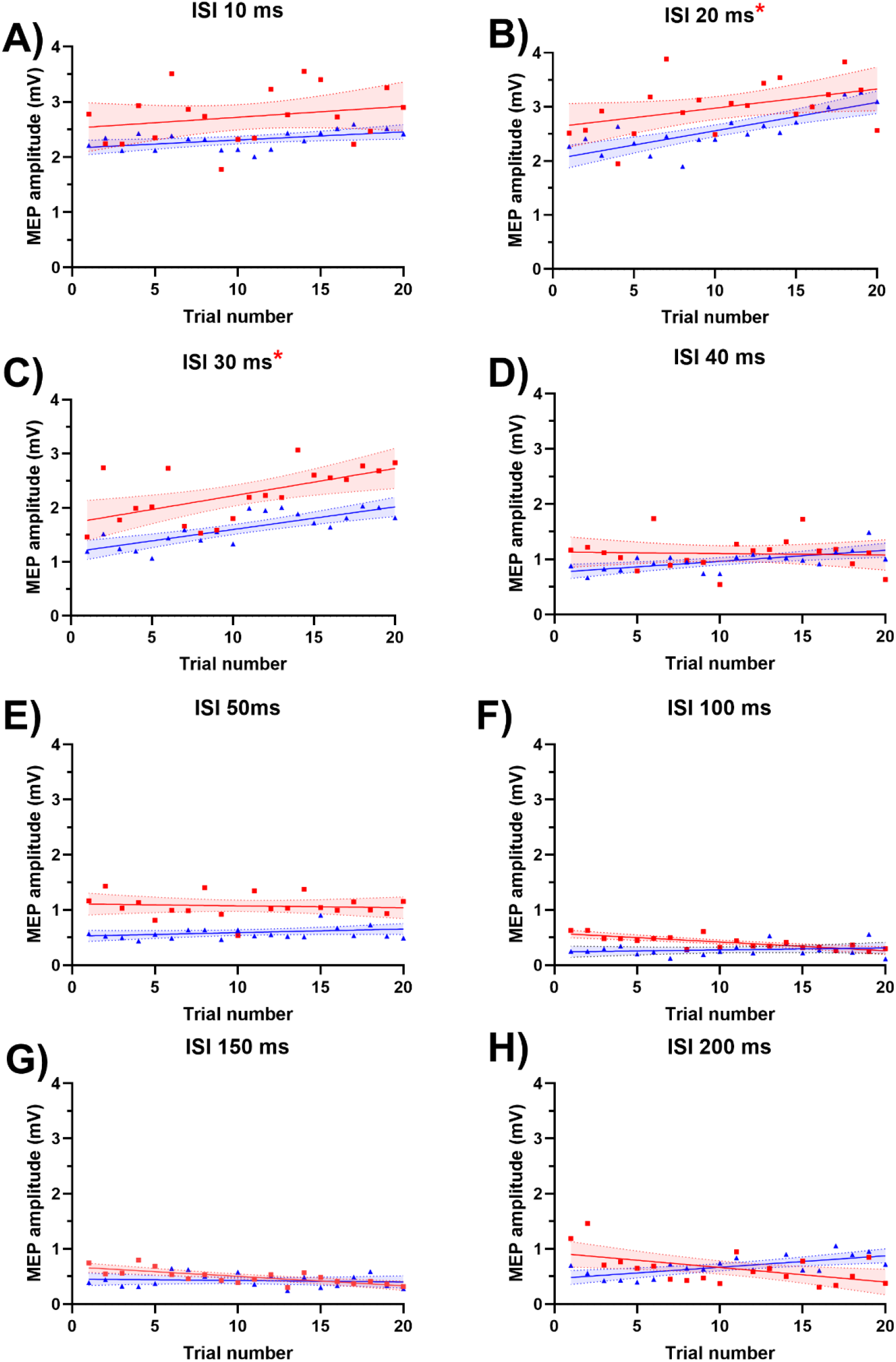
Estimated trends of changes in conditioned motor evoked potential (MEP) amplitude across trials, compared between experiment 1 (blue) and 2 (red) during eight different interstimulus intervals of paired-pulse TMS. Solid lines represent the mean regression, whereas shaded bars represent the 95% confidence intervals. The blue triangles represent the estimated marginal means of the 20 suprathreshold conditioned paired-pulse trials intermittently administered in experiment 1 and the red squares represent the those administered in experiment 2.

To better understand the findings of experiment 2, we performed post hoc analyses on this cohort. Post hoc analysis evaluating the effects of ISI in cohort 2 found significant differences in MEP amplitudes between all ISIs (*p*-value range = 0.026 to 0.0001), excluding contrasts between 50 vs 200 ms (*p* = 1.00), and 100 vs 150 ms (*p* =1.00). Post hoc analysis assessing the effects of ISI and trials within cohort 2 identified significant increase of MEP amplitudes across trials, reporting a positive slope at ISIs of 20 ms (estimated trend = 0.0356 [0.008642, 0.0626], *p* = 0.01) and 30 ms (estimated trend = 0.0527 [0.025701, 0.07974], *p* = 0.0001; figure 4). Post hoc tests contrasting slopes between ISIs showed significant differences in slope between ISIs of 20 vs 50 ms (*p* = 0.037), 20 vs 100 ms (*p* = 0.0022), 20 vs 150 ms (*p* = 0.0008), 20 vs 200 ms (*p* = 0.0212), 30 vs 50 ms (*p* = 0.0144), 30 vs 100 ms (*p* = 0.0007), 30 vs 150 ms (*p* = 0.0002) and 30 vs 200 ms (*p* = 0.0079). As with experiment 1, experiment 2 also found cumulative increases in conditioned MEP amplitudes across trials which was strongest following ISIs of 20 and 30 ms.

A separate model was performed to contrast the single pulse MEP amplitude between the two experimental cohorts. There were significant main effects of cohort (X^2^ _(1, 1032)_ = 40.83, *p* < 0.001) and trial (X^2^ _(1,1024)_ = 7.75, *p* = 0.005) on MEP amplitude. A post hoc analysis evaluating the main effect of cohort indicated that mean MEP amplitudes were significantly larger in experiment 2 compared to experiment 1 (EMD= -0.375 SE= 0.0587, *p* = <0.001) (figure 5). However, there was no evidence for a significant interaction between cohort and trial (X^2^ _(1,1022)_ = 0.3646, *p* = 0.546). Post hoc analysis found that the slope of the MEP curve significantly deviated from 0 in experiment 1 (estimated trend = 0.00734 SE= 0.0028 [0.713-1.14], *p*= 0.01), but not 2 (estimated trend = 0.005 SE= 0.003562 [1.078,1.52], *p=* 0.20). However, there was no difference in slopes between cohorts (EMD= 0.003 SE= 0.00458, *p* = 0.54). Consequently, changes in MEP amplitude across trials were comparable between cohorts (figure 5). Together, the model including both experiments suggests that cumulative changes in corticospinal excitability during suprathreshold single pulse TMS are a robust finding that replicated across two independent cohorts.

**Figure 5:**
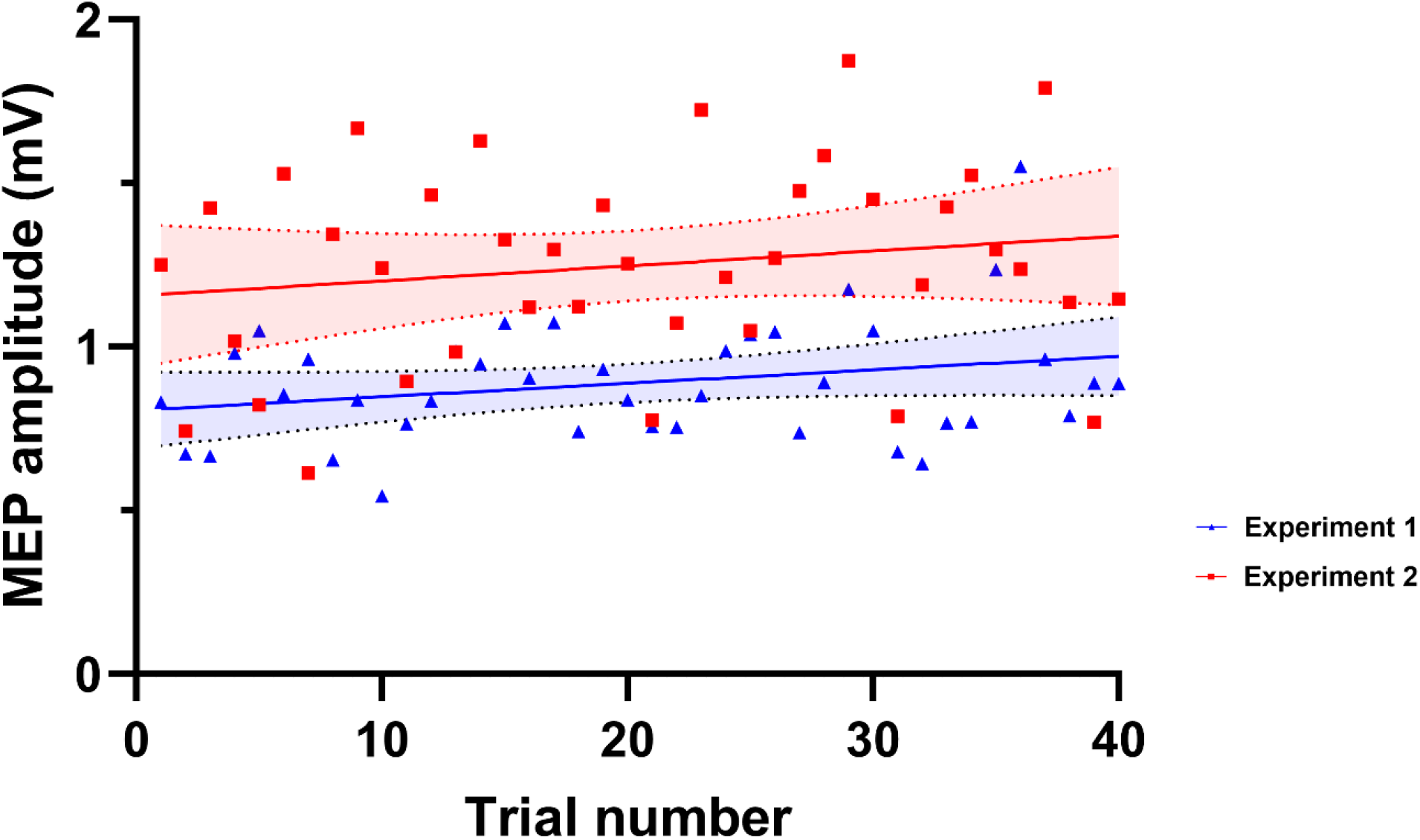
Estimated trends of changes in motor evoked potential (MEP) amplitude across trials between experiment 1 (blue) and 2 (red) during single pulse TMS. Solid lines represent the mean regression, whereas as shaded bars represent the 95% confidence intervals. The blue triangles represent the estimated marginal means from the 40 single-pulse trials intermittently administered in experiment 1, and the red squares represent those from experiment 2.

## 4. DISCUSSION

In the current study, we assessed whether suprathreshold paired-pulse TMS paradigms typically used to investigate facilitatory and inhibitory neural circuits resulted in cumulative changes in corticospinal excitability. Across two independent cohorts, we found evidence for cumulative increases in corticospinal excitability when using ISIs of 20 and 30 ms. Additionally, cumulative increases in MEP amplitude were also observed across single-pulse TMS trials. However, we could not find strong evidence for cumulative changes at ISIs targeting LICI (100-150 ms). These results indicate that caution is required when using suprathreshold paired-pulse TMS at short intervals to assess facilitatory circuits. However, our findings suggest that repetitive suprathreshold paired-pulse stimulation may provide a novel approach to induce plasticity within the corticospinal system.

Single- and paired-pulse TMS paradigms are typically used to assess the excitability of facilitatory and inhibitory circuits within the corticospinal system. For example, two suprathreshold pulses results in a period of facilitation of MEPs with ISIs between 10-40 ms, and a period of inhibition known as LICI with ISIs between 50-150 ms [14, 28]. However, an underlying assumption of these paradigms is that spacing trials between 3-6 s apart, and jittering the inter-trial interval, is sufficient to prevent stimulation from inducing cumulative changes in excitability. Several studies have challenged this assumption. Julkunen and colleagues (2012) showed that the mean amplitude of MEPs increased across 40 single-pulse trials at inter-trial intervals between 1 and 5 s, but not 10 s [20]. Pelliciarri and colleagues (2016) also reported increases in MEP amplitudes across 200 trials following single-pulse TMS at inter-trial intervals of 4 s [19]. In both studies, increases were evident regardless of whether the inter-trial interval was fixed or jittered. Similar cumulative increases in corticospinal excitability have also been reported during blocks of paired-pulse stimulation targeting excitatory circuits at short ISIs (1.5-15 ms), which have informed the development of potential plasticity boosting interventions [22, 29, 30]. However, paired-pulse paradigms targeting inhibitory circuits at short (2-3 ms) and long ISIs (100-150 ms) were not associated with cumulative changes in corticospinal excitability [22].

Our findings are broadly in line with previous work. We found increases in MEP amplitude across trials using a suprathreshold conditioning pulse and ISIs associated with facilitation of MEPs (20-30 ms) in two independent cohorts. We observed cumulative increases in MEP amplitude despite using a jittered inter-trial interval and randomising the order of different ISI conditions within stimulation blocks. These findings align with increased MEP amplitudes using a sub-threshold ICF protocol [22]. Furthermore, we also found evidence for an increase in MEP amplitude across trials following single-pulse TMS, replicating previous reports of cumulative changes in corticospinal excitability [18, 31]. In contrast, we did not find any strong evidence for cumulative changes in corticospinal excitability using ISIs associated with LICI (100-150 ms). These findings are in line with previous work, which did not find online changes in MEP amplitude using the LICI paradigm [22].

The mechanisms underlying the cumulative changes in corticospinal excitability during paired-pulse TMS remain unclear. One potential mechanism is spike-timing dependent synaptic plasticity, which is thought to underlie the cumulative facilitatory effects of other repetitive TMS paradigms [21, 32, 33]. However, the facilitation of MEPs between 20-30 ms is associated with increases in both cortical and spinal excitability. For example, suprathreshold paired-pulse TMS at an ISI of 25 ms results in increases in late I-waves supporting a role of increased cortical excitability [28]. In addition, intracortical facilitation is associated with larger H-reflexes supporting increased spinal excitability [14, 15, 34]. As a result, the changes in MEP during suprathreshold paired-pulse TMS could reflect cumulative increases in either cortical or spinal excitability, or a combination of the two. The use of other techniques more sensitive to cortical activity, like combined TMS and EEG, may help to further disentangle the relative contributions of cortical and spinal excitability to the observed effect [35, 36].

There are two main implications of our findings. First, our primary finding further challenges the assumption that an inter-trial interval of >3 s with a jitter is sufficient to prevent cumulative changes in corticospinal excitability during TMS. Our work extends previous findings on single-pulse TMS, showing that similar cumulative changes are observed with paired-pulse TMS at short ISIs. As a result, care must be taken when using suprathreshold paired-pulse TMS to assess facilitatory neural circuits due to the possibility of inducing unexpected lasting changes in corticospinal excitability. Future work assessing whether longer inter-trial intervals can circumvent these cumulative changes are required. Second, our findings raise the intriguing possibility that suprathreshold paired-pulse TMS could be a novel paradigm to induce plasticity in the corticospinal system. Other repetitive paired-pulse paradigms like I-wave periodicity rTMS also induce online changes in MEP amplitude, which can last for up to 30 mins post stimulation [21, 29, 37]. In the current study, conditioned MEP amplitude increased by ∼150% across trials. Future work assessing whether these online changes translate to offline increases in corticospinal excitability are required to assess the suitability of this method for inducing plasticity.

In conclusion, we found evidence for cumulative increases in corticospinal excitability during suprathreshold paired-pulse paradigms at facilitation-related (20-30 ms), but not LICI-related (100-150 ms) ISIs. This finding was robust and replicated across two independent datasets. Our results suggest that caution is required when using paired-pulse paradigms to assess facilitatory neural circuits due to the potential for unintended changes in corticospinal excitability. On the other hand, suprathreshold paired-pulse paradigms may offer a novel way for inducing plasticity in the corticospinal system that warrants further investigation.

## Data availability statement

Data from experiment 1 is available at:https://doi.org/10.26180/8251799.v9 Data used for statistical analyses of experiment 1 and 2 is available at:https://doi.org/10.6084/m9.figshare.30454103.v2

## Author contributions

**Suraj Suresh:** Methodology, Formal analysis, Investigation, Data Curation, Writing - Original Draft, Visualization.

**Mana Biabani:** Methodology, Formal analysis, Investigation, Data Curation, Writing - Review & Editing.

**Wei-Yeh Liao:** Formal analysis, Writing - Review & Editing.

**George Opie:** Conceptualization, Writing - Review & Editing, Supervision.

**Alex Fornito:** Writing - Review & Editing.

**Mitchell Goldsworthy:** Conceptualization, Resources, Writing - Review & Editing, Supervision, Project administration.

**Nigel Rogasch:** Conceptualization, Resources, Writing - Review & Editing, Supervision, Project administration.

